# Analysis at single-cell resolution identifies a stable mammalian tRNA-mRNA interface and increased translation efficiency in neurons

**DOI:** 10.1101/2021.06.28.450167

**Authors:** William Gao, Carlos J. Gallardo-Dodd, Claudia Kutter

## Abstract

The correlation between codon and anticodon pools influences the efficiency of translation, but whether differences exist in these pools across individual cells is unknown. We determined that codon usage and amino acid demand are highly stable across different cell types using single-cell RNA-sequencing atlases of adult mouse and fetal human. After demonstrating the robustness of ATAC-sequencing for analysis of tRNA gene usage, we quantified anticodon usage and amino acid supply in adult mouse and fetal human single-cell ATAC-seq atlases. We found that tRNA gene usage is overall coordinated across cell types, except in neurons which clustered separately from other cell types. Integration of these datasets revealed a strong and statistically significant correlation between amino acid supply and demand across almost all cell types. Neurons have an enhanced translation efficiency over other cell types, driven by an increased supply of tRNA^Ala^ (AGC) anticodons. This results in faster decoding of the Ala-GCC codon, as determined by cell-type specific ribosome profiling, and a reduction of tRNA^Ala^ (AGC) anticodon pools may be implicated in neurological pathologies. This study, the first such examination of codon usage, anticodon usage, and translation efficiency at single-cell resolution, identifies conserved features of translation elongation across mammalian cellular diversity and evolution.

## Introduction

During translation elongation, the ribosome moves three nucleotides at a time along the mRNA transcript, as each codon complementarily binds to a corresponding anticodon triplet on a tRNA, which is charged with a specific amino acid that is then added to the growing polypeptide chain (Dever et al. 2018). Since translation elongation relies on the codon-anticodon interaction, matching or mismatching of codon and anticodon pools may influence the elongation rate. Indeed, it has been shown in bacteria such as *Escherichia coli* and unicellular eukaryotes such as *Saccharomyces cerevisiae* that matching of these pools results in increased translation efficiency, leading to higher production of proteins. Conversely, mismatching of these pools yields lower protein production (Quax et al. 2015).

Codon and anticodon pools have also been measured in multicellular organisms, such as mammals, which pose several additional challenges. In particular, the mRNA and tRNA levels may differ across tissues and cell types and quantitation can be biased because of experimental challenges (Wong et al. 2012). The development of bulk next-generation sequencing (Stark et al. 2019) in the past two decades has allowed for examination of these mRNA and tRNA pools across a few mammalian tissues, revealing some tissue-specific differences but an overall stability in both codon and anticodon usage (Dittmar et al. 2006; Schmitt et al. 2014; Rak et al. 2018; Pinkard et al. 2020). In these studies, codon pools were measured primarily using bulk RNA-sequencing. However, due to several special features of tRNA genes, anticodon pools have been measured with a variety of RNA- and DNA-based methods that are subject to different biases. For example, RNA-based methods for tRNA gene expression include microarray (Chou et al. 2004; Dittmar et al. 2006) and next generation sequencing approaches (Zheng et al. 2015; Gogakos et al. 2017; Shigematsu et al. 2017; Xu et al. 2019; Kugelberg et al. 2021). These methods must overcome the challenge of producing complementary DNA from tRNAs which are not only highly structured but also extensively modified (Rak et al. 2018; Suzuki 2021). As a result, no “gold standard” approach exists to measure tRNA abundances, with a recent study showing that the tRNA levels obtained from different methods are lowly correlated (ρ ~ 0.22-0.62) (Pinkard et al. 2020). Alternatively, Chromatin Immunoprecipitation with massively parallel DNA sequencing (ChIP-seq) can be performed in bulk with an antibody targeting an active subunit of RNA Polymerase III (Pol III) that transcribes tRNA genes (Dieci et al. 2007). Examining active Pol III occupancy on DNA quantifies the amount of tRNA transcription and is not prone to biases resulting from tRNA structure and modification (White 2011). Moreover, because ChIP-seq reads often extend past the exact sequence of the tRNA gene, intergenic DNA can be used to resolve tRNA genes with otherwise identical sequences. However, Pol III ChIP-seq only probes pre-tRNA transcription. Pre-tRNAs undergo several more steps before becoming ready-to-translate tRNAs (Wolin and Matera 1999; Phizicky and Hopper 2010).

Using these methods of quantifying codon and anticodon usage, previous studies have examined how correlated these pools are, using this as a proxy for the efficiency of translation elongation (often referred to as “translation efficiency”). Analysis of codon and anticodon pools across the tree of life has revealed several unifying features, such as the necessity of wobble base pairing between codons and anticodons. Indeed, all organisms use fewer than 61 anticodons to decode the 61 sense codons because decoding can occur with wobble interactions between the first anticodon position (tRNA nucleotide 34) and the third codon position (Rak et al. 2018). In particular, the most well-characterized modifications include G:U and adenosine-to-inosine 34 (A34-to-I) wobbling. The latter, performed by adenosine deaminases (ADATs), expands the repertoire of ANN anticodons to decode not only NNU codons but also NNC and NNA codons (Torres et al. 2014).

In unicellular organisms such as *E. coli* and *S. cerevisiae*, the most frequent codons also correspond to the most abundant anticodons and this explains a great deal of patterns in synonymous codon usage bias (Rocha 2004). In contrast, work in mammals has suggested that adaptation to anticodon pools cannot explain the majority of synonymous codon usage, which appears to be more strongly influenced by mutational biases and drift (dos Reis et al. 2004), in particular GC-biased gene conversion (Pouyet et al. 2017a). Despite weak correlations between synonymous codon usage bias and anticodon levels (Novoa and Ribas de Pouplana 2012), the overall correlation between codon and anticodon pools in mammals are nonetheless quite strong and stable (Rudolph et al. 2016). Yet all of these observations in mammals have been made using bulk sequencing.

However, a major limitation of using bulk methods to quantify codon and anticodon pools is that by aggregating data from multiple different cell types within a tissue, they may blur out heterogeneity across cell types. Since cell types use not only different levels of the same proteins but also different types of proteins to perform their various functions, it is possible that their codon pools are different. Additionally, the tRNA gene usage across cell types may also differ to match their codon pools (Dittmar et al. 2006; Rak et al. 2018).

The advent of multi-omics single-cell technologies, and the production of publicly available whole organism single-cell atlases allows us to examine codon and anticodon usage at cell-type specificity and uncover heterogeneity that cannot be detected in bulk. Here, we used an adult mouse (Schaum et al. 2018) and fetal human scRNA-seq atlas (Cao et al. 2020) to examine codon usage and identified strong similarities across cell types. We then demonstrated the reliability of using scATAC-seq for quantification of anticodon usage and examined anticodon usage in corresponding adult mouse (Cusanovich et al. 2018) and fetal human (Domcke et al. 2020) scATAC-seq datasets. We uncovered stability in anticodon usage across cell types, but with noticeable differences in neurons. Finally, we calculated translation efficiency by correlating amino acid (AA) demand and supply from the mRNA and tRNA side, respectively. We found a strong, statistically significant correlation in these pools across all cell types, but with enhanced translation efficiency in neurons. By examining these important contributors to translation elongation for the first time at the single-cell level, this study paves the way for future high-resolution studies of mammalian translation.

## Results

### Codon usage is highly similar across cell types

To investigate whether codon usage differs across mammalian cell types, we analyzed single-cell RNA-sequencing (scRNA-seq) atlases from adult mouse (AM) and fetal human (FH). The AM and FH datasets consist of cells from 20 and 15 tissues, respectively (**Figs. 1A-B, S1A, Table S1**). Cell type annotations have been provided for all individual cells that passed quality filters.

**Fig. 1:**
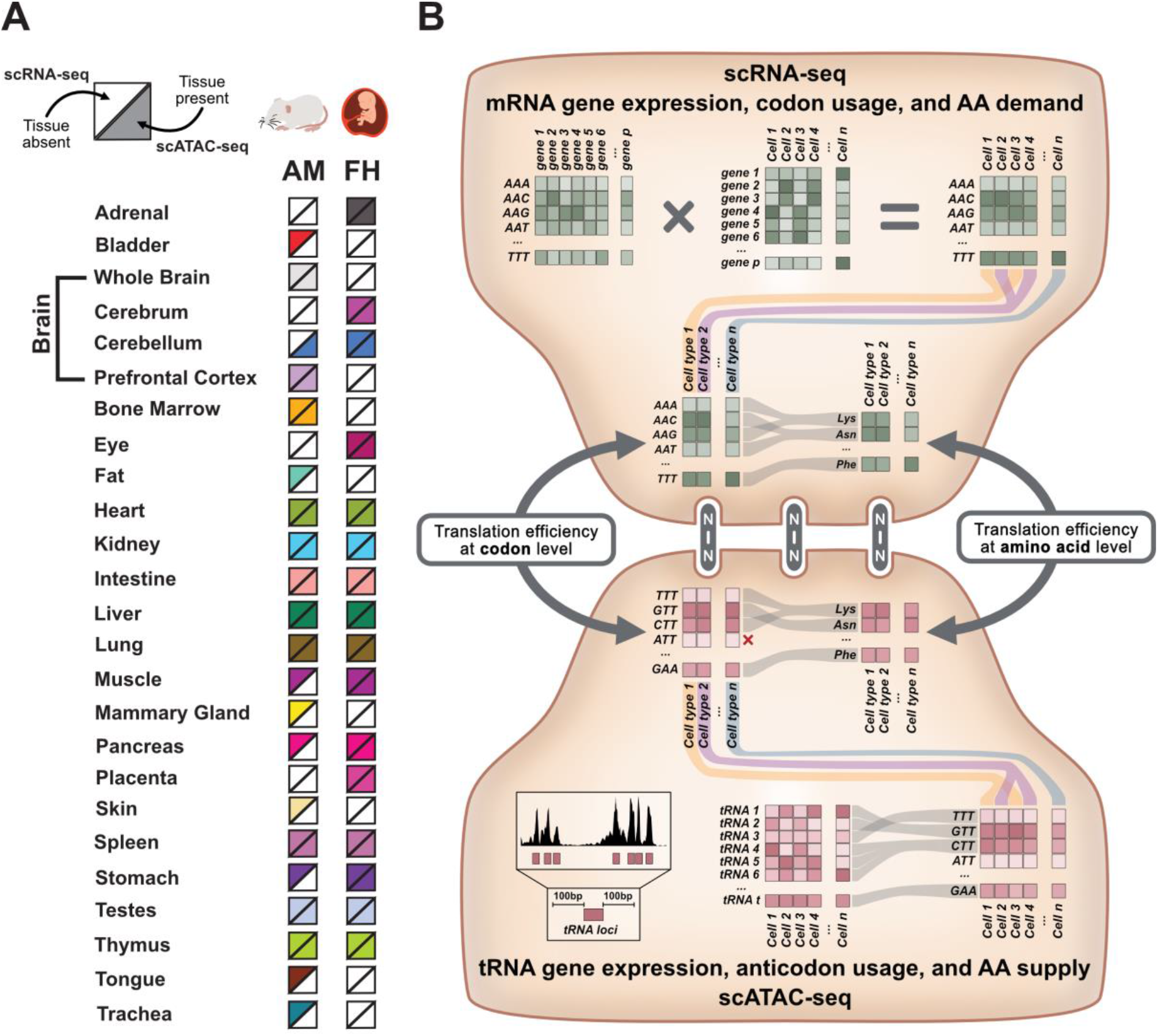
Overview of the approach for single-cell analysis of translation efficiency. **(A)** To examine a diverse set of cell types across multiple tissues, scRNA-seq and scATAC-seq atlases produced for adult mouse (AM) and fetal human (FH) were analyzed. For each organism, each square is color-filled if scRNA-seq (top half) or scATAC-seq data (bottom half) are present. **(B)** top: mRNA gene expression, codon usage, and amino acid demand can be quantified using single-cell RNA-sequencing (scRNA-seq). Each gene’s sense codon frequencies are weighted by gene expression counts. Codon usage from individual cells is pooled at the cell-type level and can be combined based on amino acid demand. bottom: tRNA gene expression, anticodon usage, and AA supply can be quantified using the single-cell Assay for Transposase Accessible Chromatin (scATAC-seq). First, the Tn5 transposase insertions mapping within 100 nucleotides of tRNA genes are quantified, creating a tRNA gene expression matrix. Cells are pooled at the cell-type level and can be combined into anticodon isoacceptor families and amino acid isotypes. center: scRNA-seq and scATAC-seq data can be integrated to calculate translation efficiency. For more detailed description, see Methods section.

Different cell types express a diverse assembly of genes at dissimilar levels, and the high expression of specific marker genes is often used for cell type annotation. When pooling the gene expression profiles from individual cells of the same cell type, we observed the expected clustering of cell types based on function and tissue origin (**Fig. 2A, Tables S2, S9**). For example, endothelial and stromal cells from several organs clustered together, while in other cases, cell types from the same tissue clustered together due to expression of tissue-specific genes, as observed previously (Schaum et al. 2018). Clustering of cell types by origin and function was less defined in the FH dataset, perhaps due to the collection of these samples in early development (72-129 days) (Cao et al. 2020).

**Fig. 2:**
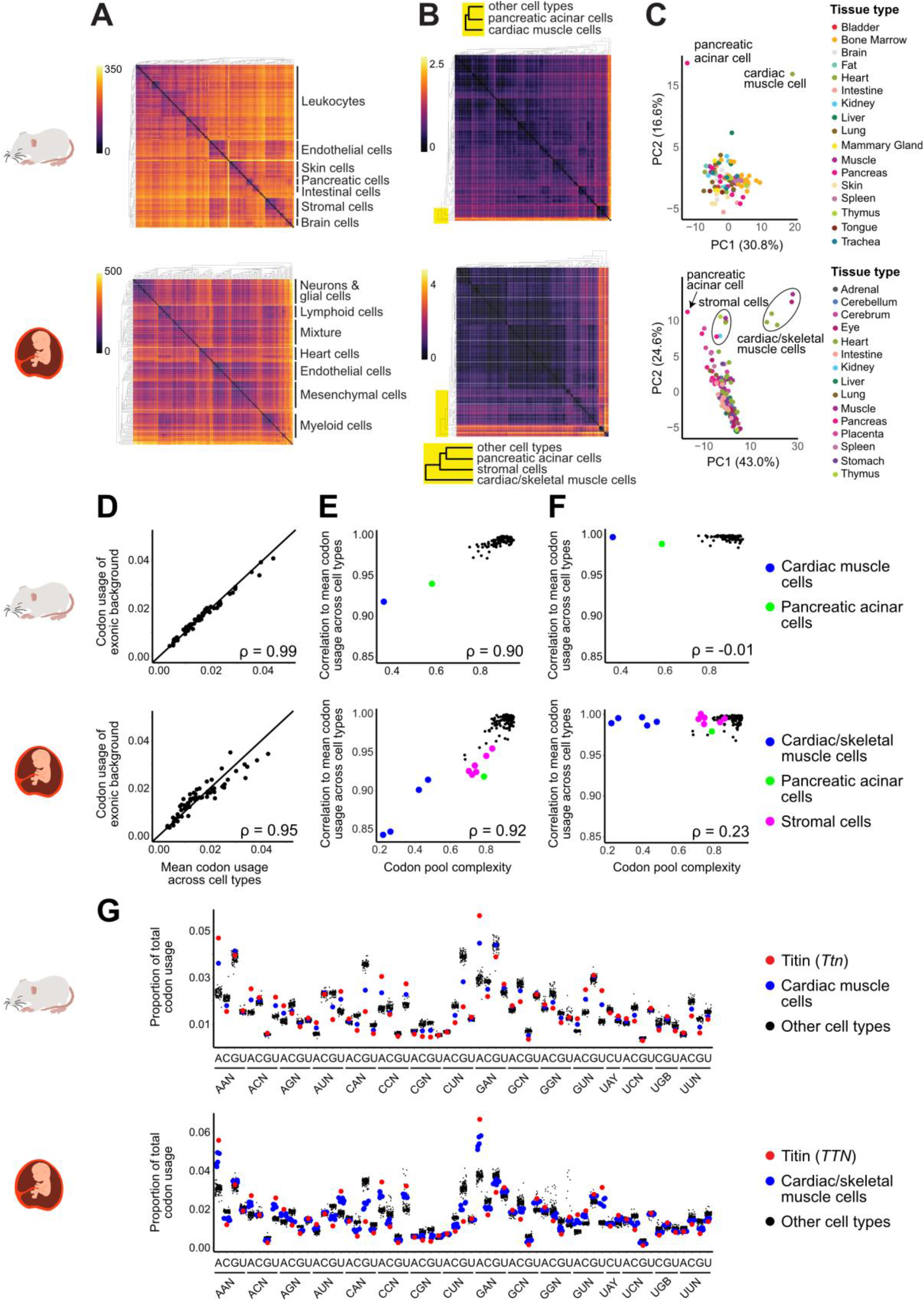
Codon usage is highly stable across cell types, with outliers driven by reduced codon pool complexity. **(A-B)** Heatmaps showing Euclidean distance in (A) gene expression and (B) codon usage across cell types for AM (top) and FH (bottom) **(C)** Principal component analysis (PCA) plots show codon usage across cell types. **(D-F)** Scatter plots demonstrate correlation of (D) mean codon usage across all cell types to the exonic background (unweighted genome-wide codon usage). Each point corresponds to one of the 61 sense codons, (E) each cell type’s codon pool complexity (percentage of the total codon pool contributed by the top 10 genes) to the correlation of its cell type to the mean codon usage across all cell types and (F) same quantities as (E), but with each cell type’s codon usage calculated while ignoring the codon contribution of the top 10 codon pool contributing genes. **(G)** Jitter plots show the proportion of total codon usage for each of the 61 sense codons in each cell type. Titin (red) drives the outlier status of cardiac and skeletal muscle cells (blue). IUPAC nucleotide codes are used in the x-axis (B = A, C, U; R = A, G; Y = C, U).

To calculate the codon usage across cell types, we weighted the 61 sense codon frequencies of each protein-coding gene by their cell-type-specific expression (**Fig. 1B** top, **Tables S3-S4, S10-S11**). The sense codon frequencies were determined from the trinucleotide frequencies of the coding sequences (CDS) of each protein-coding gene. The longest open reading frame (ORF) per protein-coding gene was used to determine its codon frequencies, as isoform information is not available for all cell types in the scRNA-seq datasets but remains an active area of research (Uhlén et al. 2015).

In contrast to gene expression, we found a strong similarity in codon usage across cell types. The Euclidean distances between codon usages across almost all cell types is small (**Fig. 2B**) and a Principal Components Analysis (PCA) plot of codon usages across cell types reveals that nearly all of them colocalize, without distinct clustering (**Fig. 2C**). Previous analysis of codon usage in bulk had also suggested highly stable codon pools across tissues (Schmitt et al. 2014), but it remained unknown whether particular cell types had distinct codon usage profiles that were undetected in aggregate. Here, we found that codon usage was highly similar across the majority of cell types. Moreover, the mean codon usage across all cell types was highly similar to the exonic background (the overall genome codon frequencies unscaled by gene expression) (**Fig. 2D**), as established in bulk (Schmitt et al. 2014). In other words, our analysis at the cell-type level largely recapitulated findings from bulk analysis due to true homogeneity in codon usage across cell types, rather than averaging of heterogeneous codon usages.

### Cell types that deviate from codon usage have low codon pool complexity

While most cell types have similar codon usages, some cell types were distinct. In both AM and FH, cardiac muscle cells and pancreatic acinar cells had larger Euclidean distances from other cell types and resided away from the main cluster in PCA (**Figs. 2B-C**). Stromal cells from some FH tissues also segregated from the main cluster, while others did not (**Fig. 2C**). No AM stromal cell populations were outside the main cluster. Thus, the differential codon usage in some fetal stromal cell types may be present only in early development. Skeletal muscle cells, which were not present in the AM dataset, clustered in codon usage with cardiac muscle cells in the FH dataset (**Fig. 2C**).

Since codon usage was computed by weighting each gene’s codon frequencies by its expression, differential codon usage in cardiac/skeletal muscle cells and pancreatic acinar cells could be influenced by the diversity of protein-coding genes that contribute to their codon pools. A cell type could be an outlier in codon usage because of widespread differences in codon usage across numerous expressed genes, or because its codon usage is skewed by a few highly expressed genes with large codon demands. To determine which was the case, we calculated each gene’s codon pool contribution per cell type, defined as the number of codons for that gene multiplied by its cell-type-specific expression. Thus, longer genes that are also highly expressed will have the largest codon pool contributions. We define codon pool complexity as the proportion of the codon pool that is not contributed by the top N genes, ranked by codon pool contribution. Thus, a low codon pool complexity means that the top N genes’ codon pool contributions are a large percentage of the total codon pool, while a high codon complexity indicates that many genes contribute a small amount to the codon pool.

We found that cell types with low codon pool complexity (calculated for several values of N ranging from 5 to 100) tend to be those that are outliers in codon usage, with the lowest correlations in codon usage to the mean codon usage across all cell types (**Fig. 2E**). By ignoring the codon contribution of the top 10 genes, their codon usages become indistinguishable from the other cell types (**Fig. 2F**). Therefore, these cell types were outliers in codon usage because relatively few protein-coding genes with skewed codon frequencies comprised the bulk of their codon pools. For example, cardiac and skeletal muscle cells express high levels of *TTN*, the gene encoding titin. It is the largest known protein (27,000-35,000 amino acids, depending on the isoform) and important for the passive elasticity of muscle (Lewinter and Granzier 2010). Due to its remarkable length and high expression in cardiac and skeletal muscle cells, this gene accounts for between 48-71% of their codon pools (56% in AM cardiac muscle cells, 48-71% in FH cardiac/skeletal muscle cells). Thus, when titin’s usage of particular codons differs from the mean codon usage across cell types, cardiac and skeletal muscle cells become outliers for those codons (**Fig. 2G**).

### scATAC-seq is robust for measuring tRNA gene usage

Having established that codon usage is highly stable across most cell types, we next investigated whether anticodon usage would also be similar. Previous work has examined tRNA gene usage in different cell lines and in tissues, identifying tissue-specific differences in tRNA gene usage (Dittmar et al. 2006; Pinkard et al. 2020; Schmitt et al. 2014; Rudolph et al. 2016). Though existing RNA- and DNA-based tRNA quantification methods have their respective advantages and disadvantages, none have been adapted for analysis in single cells. To address this major limitation, we investigated whether the Assay for Transposase-Accessible Chromatin (ATAC-seq) could be used to examine tRNA gene usage. This assay, uses a hyperactive Tn5 transposase that cuts at open chromatin (Buenrostro et al. 2015). Thus, loci enriched for Tn5 insertions correspond to regions of open chromatin, at which transcription factors and RNA Pol can bind. Indeed, chromatin accessibility in gene bodies often correlates well with transcription levels (Klemm et al. 2019). Furthermore, several different methods have been developed for single-cell analysis (Pott and Lieb 2015; Baek and Lee 2020),

Because scATAC-seq has not been used before for analyzing tRNA genes, we assessed its robustness by determining whether it is internally consistent (different scATAC-seq pipelines produce similar quantifications of tRNA accessibility), externally consistent (concordant with expectation), and of adequate resolution (can distinguish between tRNA genes located in tight genomic clusters). We found that tRNA gene accessibility from three available scATAC-seq of AM brain (aggregated in pseudobulk) (Cusanovich et al. 2018; Lareau et al. 2019) were strongly correlated at the tRNA gene, anticodon isoacceptor, and amino acid (AA) isotype levels (**Figs. S1, S2**), demonstrating the internal consistency of ATAC-seq for tRNA gene quantification. These data were also highly correlated at all three levels with a bulk AM brain ATAC-seq dataset (Liu et al. 2019) and, importantly, with tRNA abundances measured by Pol III ChIP-seq, the established DNA-based tRNA quantification method. This suggests that the chromatin accessibility of tRNA genes is a suitable measure for their expression (**Fig. 3A**). The correlations across these different datasets were consistently high (Spearman’s rank correlation coefficients, ρ ~ 0.7 to 0.9), which is much higher than the ones observed when comparing across different RNA-based quantification methods (Pearson’s correlation coefficients, r ~ 0.2 to 0.6) (Pinkard et al. 2020). To validate ATAC-seq irrespective of other approaches, we also examined whether highly confident functional tRNA genes were more accessible than low-confidence tRNA gene predictions (Lowe and Chan 2016). We found that high-confidence tRNA genes were generally much more accessible than low-confidence ones (**Fig. 3B**). Finally, since tRNA genes tend to be genomically organized in dense clusters, we assessed whether scATAC-seq data provided enough resolution to differentiate accessibility between very close tRNA genes. Indeed, scATAC-seq displays individual peaks for tRNA genes in clusters in which tRNA genes were fewer than 300 nucleotides apart (**Fig. 3C**).

**Fig. 3:**
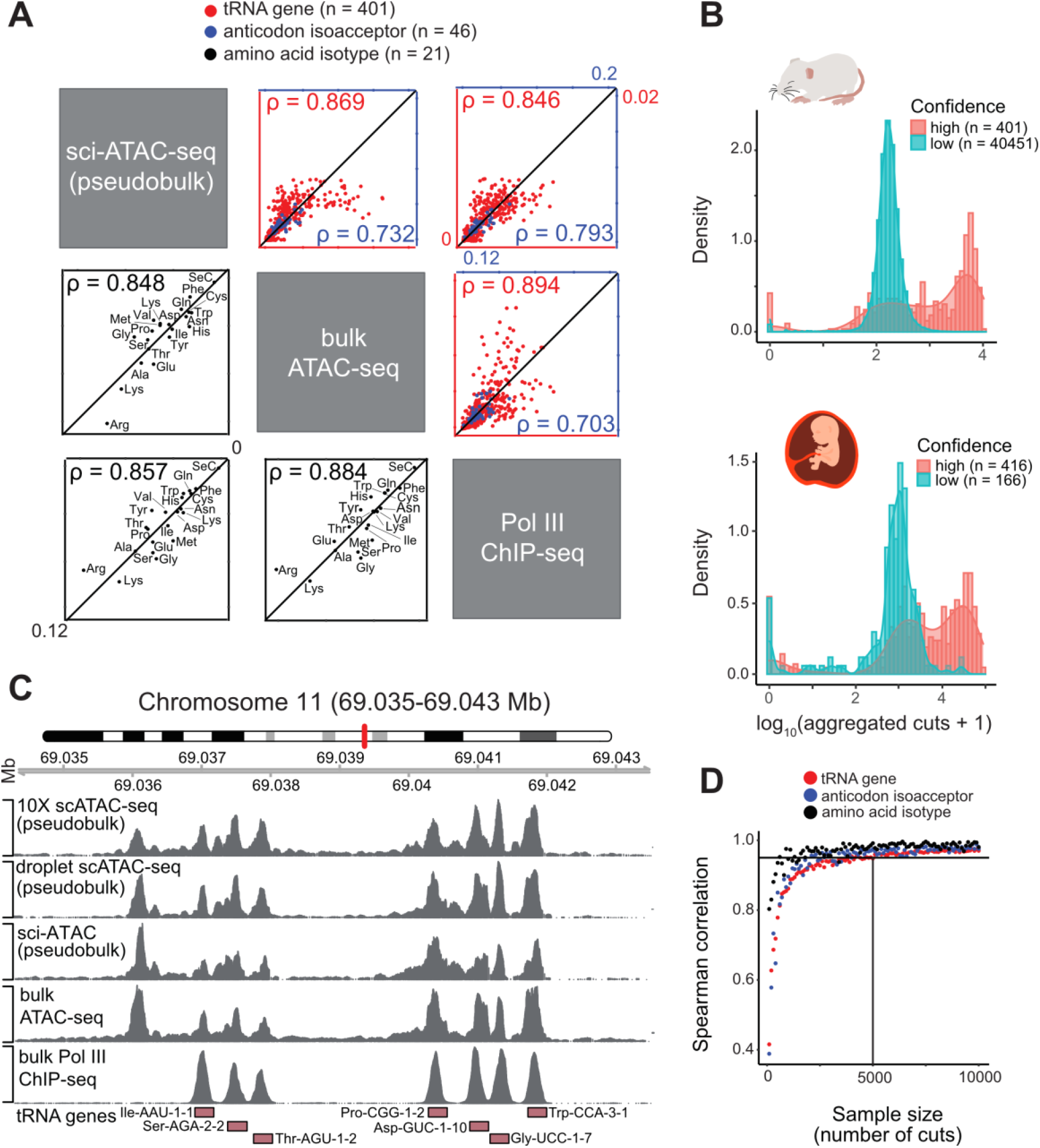
Single-cell ATAC-sequencing is robust for measuring tRNA gene usage. **(A)** Scatter plots correlates AM brain scATAC-seq (aggregated in pseudobulk), bulk ATAC-seq, and Pol III ChIP-seq dataset on the tRNA gene (red), anticodon isoacceptor (blue) and amino acid isotypes (black) level. Spearman’s rank correlation coefficients are indicated on the bottom left and top right corners. **(B)** Density plots (in log-scale) show the total number of cuts from aggregated scATAC-seq data from the AM and FH scATAC-seq atlases, based on confidence predicted from tRNAscan-SE. **(C)** Genome browser view illustrates a mouse tRNA gene cluster on chromosome 11. The location of tRNA genes is shown at the bottom, including upstream and downstream 100 nt flanking regions. Different tRNA genes, including those as close as 220 nucleotides apart (*Asp-GUC-1-10* and *Gly-UCC-1-7*), have distinct peaks in three scATAC-seq datasets (pseudobulk), a bulk ATAC-seq dataset, and a Pol ChIP-seq dataset (all AM brain). The leftmost peak present in all ATAC-seq datasets but absent in the Pol III ChIP-seq dataset corresponds to the promoter of protein-coding gene *Ctc1*. This peak is ignored when quantifying tRNA gene usage because it falls outside of 100 nts of a tRNA gene. **(D)** Scatter plot determines how many total cuts are needed after pooling cells of the same cell type to obtain reliable information of tRNA gene, anticodon isoacceptor, and amino acid isotype usage. A sample size of 5,000 cuts per cell type consistently yielded a Spearman’s rank correlation coefficient greater than 0.95 to the aggregated scATAC-seq data from AM and FH.

### scATAC-seq is robust for measuring tRNA gene usage

Having established that codon usage is highly stable across most cell types, we next investigated whether anticodon usage would also be similar. Previous work has examined tRNA gene usage in different cell lines and in tissues, identifying tissue-specific differences in tRNA gene usage (Dittmar et al. 2006; Pinkard et al. 2020; Schmitt et al. 2014; Rudolph et al. 2016). Though existing RNA- and DNA-based tRNA quantification methods have their respective advantages and disadvantages, none have been adapted for analysis in single cells. To address this major limitation, we investigated whether the Assay for Transposase-Accessible Chromatin (ATAC-seq) could be used to examine tRNA gene usage. This assay, uses a hyperactive Tn5 transposase that cuts at open chromatin (Buenrostro et al. 2015). Thus, loci enriched for Tn5 insertions correspond to regions of open chromatin, at which transcription factors and RNA Pol can bind. Indeed, chromatin accessibility in gene bodies often correlates well with transcription levels (Klemm et al. 2019). Furthermore, several different methods have been developed for single-cell analysis (Pott and Lieb 2015; Baek and Lee 2020),

Because scATAC-seq has not been used before for analyzing tRNA genes, we assessed its robustness by determining whether it is internally consistent (different scATAC-seq pipelines produce similar quantifications of tRNA accessibility), externally consistent (concordant with expectation), and of adequate resolution (can distinguish between tRNA genes located in tight genomic clusters). We found that tRNA gene accessibility from three available scATAC-seq of AM brain (aggregated in pseudobulk) (Cusanovich et al. 2018; Lareau et al. 2019) were strongly correlated at the tRNA gene, anticodon isoacceptor, and amino acid (AA) isotype levels (**Figs. S1, S2, Tables S5-S7, S12-14**), demonstrating the internal consistency of ATAC-seq for tRNA gene quantification. These data were also highly correlated at all three levels with a bulk AM brain ATAC-seq dataset (Liu et al. 2019) and, importantly, with tRNA abundances measured by Pol III ChIP-seq, the established DNA-based tRNA quantification method. This suggests that the chromatin accessibility of tRNA genes is a suitable measure for their expression (**Fig. 3A**). The correlations across these different datasets were consistently high (Spearman’s rank correlation coefficients, ρ ~ 0.7 to 0.9), which is much higher than the ones observed when comparing across different RNA-based quantification methods (Pearson’s correlation coefficients, r ~ 0.2 to 0.6) (Pinkard et al. 2020). To validate ATAC-seq irrespective of other approaches, we also examined whether highly confident functional tRNA genes were more accessible than low-confidence tRNA gene predictions (Lowe and Chan 2016). We found that high-confidence tRNA genes were generally much more accessible than low-confidence ones (**Fig. 3B**). Finally, since tRNA genes tend to be genomically organized in dense clusters, we assessed whether scATAC-seq data provided enough resolution to differentiate accessibility between very close tRNA genes. Indeed, scATAC-seq displays individual peaks for tRNA genes in clusters in which tRNA genes were fewer than 300 nucleotides apart (**Fig. 3C**).

While this demonstrates the robustness of scATAC-seq for quantifying tRNA gene usage, a practical concern was that scATAC-seq data could be very sparse since there are normally only two copies of DNA per cell (Baek and Lee 2020). Moreover, since tRNA genes are very short and occupy a small percentage of mammalian genomes, the number of cuts to tRNA genes per cell could be low at normal sequencing depths (**Fig. S1B**). To determine how many cuts to tRNA genes (gene body plus 100 nts upstream and downstream, as typically done for Pol III ChIP-seq data (Kutter et al. 2011)) are required to obtain reliable measurements, we took different sample sizes of cuts from pseudobulked scATAC-seq data and measured the correlation of these samples to the overall tRNA gene usage (pseudobulked scATAC-seq data without downsampling). At small sample sizes, the correlations at the tRNA gene, anticodon, and AA supply levels were all quite low and highly variable, while a sample size of greater than 5,000 cuts reliably yielded high correlations (ρ > 0.95) at all three levels (**Fig. 3D**). Thus, we analyzed cell-type specific tRNA gene usage by pooling all cells annotated as belonging to a particular cell type. Only cell types with more than 5,000 total cuts after pooling were considered reliable for tRNA gene expression analysis.

### Neuronal cell types are outliers from an otherwise stable anticodon pool

To examine tRNA gene expression, anticodon usage, and AA supply across many different cell types, we analyzed scATAC-seq atlases that corresponded to the scRNA-seq atlases (**Figs. 1, S1**). The AM and FH scATAC-seq atlas contained 13 and 15 tissues, respectively (Cusanovich et al. 2018; Domcke et al. 2020). For the cell types remaining after filtering for sufficient scATAC-seq cuts, we combined the expressions of tRNA genes on the anticodon level and computed their usage across cell types (**Fig. 1B**, bottom). We noticed that while anticodon usage was generally more variable than codon usage, it was still similar across cell types, as indicated by similar Euclidean distances in anticodon usage across most cell types (**Fig. 4A**) and the presence of a large PCA cluster corresponding to most cell types (**Fig. 4B**). However, we also observed a distinct neuron-specific cluster in AM. This brain neuron cluster was also present in FH, although less pronounced, again possibly due to decreased cell differentiation in early development. Additionally, the FH dataset also included other neuronal cell types from the eye and enteric nervous system absent from the AM dataset as well as neuroendocrine cells.

**Fig. 4:**
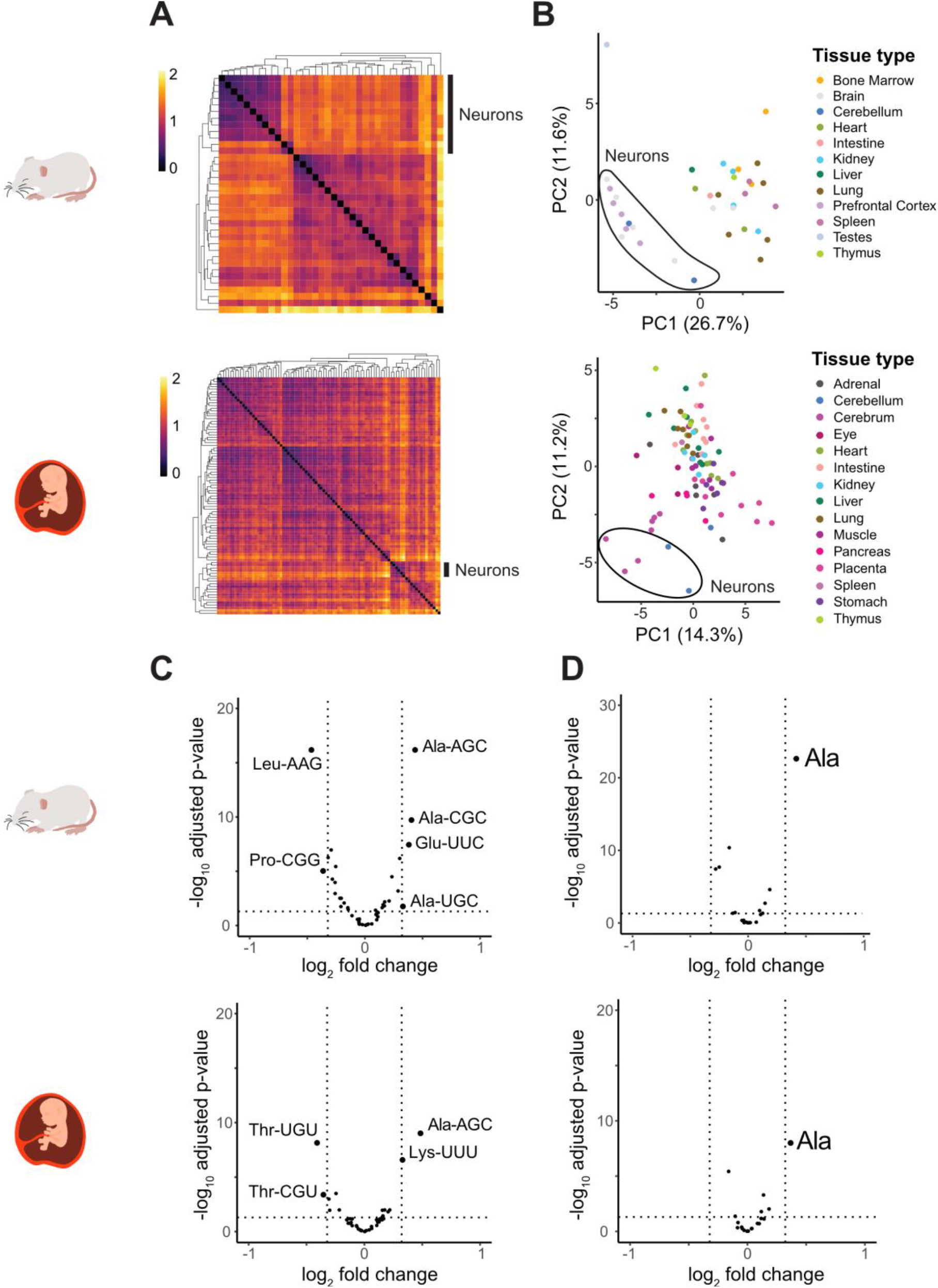
Anticodon usage is also similar across cell types, with brain neurons clustering separately. **(A)** Heatmaps show the Euclidean distance between anticodon usage across cell types. Only cell types with more than 5,000 scATAC-seq cuts were analyzed (see Fig. 3D). The neuronal cluster is indicated with a black line on the right side of the heatmap. **(B)** PCA plots separate of anticodon usage across cell types, with brain neuron cluster indicated. **(C-D)** Volcano plots of (C) anticodon usage and (D) amino acid supply between brain neurons and all other cell types, displaying −log_10_ adjusted p-values and log_2_ fold change (FC), as determined using DESeq2. Vertical lines indicate a fold change greater than 25% in one direction. All figures: top (AM), bottom (FH).

To examine what caused brain neurons to cluster separately from other cell types, we performed differential analysis at the anticodon level. We found that in both mouse and human, the anticodon AGC charged with alanine, tRNA^Ala^ (AGC), is enriched by more than 25% in brain neurons when compared against all other cell types (**Fig. S3**). In the AM brain, the two other alanine anticodon isoacceptors, tRNA^Ala^ (UCG) and tRNA^Ala^ (CCG), were also enriched in glial cells as well (**Fig. 4C**). The fourth alanine anticodon isoacceptors tRNA^Ala^ (GCG) is absent in either mammalian genome. Accordingly, when we quantified on the AA isotype level, we confirmed tRNA^Ala^ enrichment in neurons (**Fig. 4D**). Although the tRNA anticodon pools of all cell types was fairly stable, the slight deviation of one AA isotype could have significant effects on global translation (Kapur et al. 2017).

To verify that Ala isotype enrichment was observed in other datasets, we also analyzed available bulk datasets. We found that Ala supply is enriched in embryonic brain in a bulk ATAC-seq dataset of mouse embryonic development (Gorkin et al. 2020) (**Fig. S4**), a bulk ChIP-seq dataset of mouse brain and liver from early development to adulthood (Schmitt et al. 2014) (**Fig. S5**), and a RNA-based multi-tissue QuantM-tRNA-seq dataset of adult mouse (Pinkard et al. 2020) (**Fig. S6**). This, in addition to the comparison to Pol III ChIP-seq described above **(Fig. 3A**), underscores the concordance of scATAC-seq-based tRNA quantification to other methods.

### Several tRNA anticodons are enriched in neurons

While tRNA genes encoding for the same anticodon and AA isotype are often considered functionally redundant, previous work has shown that the expression of individual tRNA genes can vary quite drastically across tissues (Schmitt et al. 2014; Pinkard et al. 2020). In contrast, anticodon isoacceptor usage and AA supply are much more similar across tissues, suggesting an unknown buffering mechanism that coordinates tRNA gene expression to yield a stable supply of anticodons. This apparent buffering is also observed at the cell-type level in our analysis (**Figs. 4C-D**). However, the assumption that tRNA genes of the same anticodon isoacceptor family are functionally identical may be untrue (Pan 2018).

Therefore, we also examined differential expression at the individual tRNA gene level. As with anticodon usage, PCA at the tRNA gene level for both datasets indicated a brain neuron cluster (**Fig. S7**). By performing a differential gene analysis comparing brain neurons against all other cells in the AM and FH datasets, we observed that several tRNA genes that are enriched in neurons have not only the same anticodon but are also syntenic in the human and mouse genomes (**Figs. 5A-B**). These syntenic, neuronally-enriched tRNA genes include *Arg-TCT-4-1* (*n-Tr20*), which is one of the few tRNA genes that has been well-characterized. A tRNA maturation-inhibiting mutation in *Arg-UCU-4-*1 combined with deletion of a ribosome recycling factor causes ribosome stalling at cognate Arg-AGA codons and leads to ataxia and early death in mice (Ishimura et al. 2014). Further work has demonstrated that loss of this tRNA reduces seizure susceptibility (Kapur et al. 2020). We found two other syntenic, neuronally-enriched tRNA genes *Ala-AGC-3-1* (same name in both mm10 and hg19) and *Ile-UAU-2-1* (in mm10) corresponding to *Ile-UAU-2-3* (in hg19). Due to the cell-type resolution of scATAC-seq, we observed a clear difference in expression of these tRNA genes in neurons compared to glial cells. Additionally, in the FH dataset, which contained cells for the stomach and adrenal gland that are absent from the AM dataset, we found high expression of neuronally-enriched tRNA genes in non-brain neurons, including neurons in the eye (ganglion cells) and the enteric nervous system, as well as in neuroendocrine cells (chromaffin cells, sympathoblasts, Schwann cells and islet endocrine cells) (**Fig. 5B**). In summary, a specific subset of tRNA genes is uniquely transcribed in cells of the neuronal lineage.

**Fig. 5:**
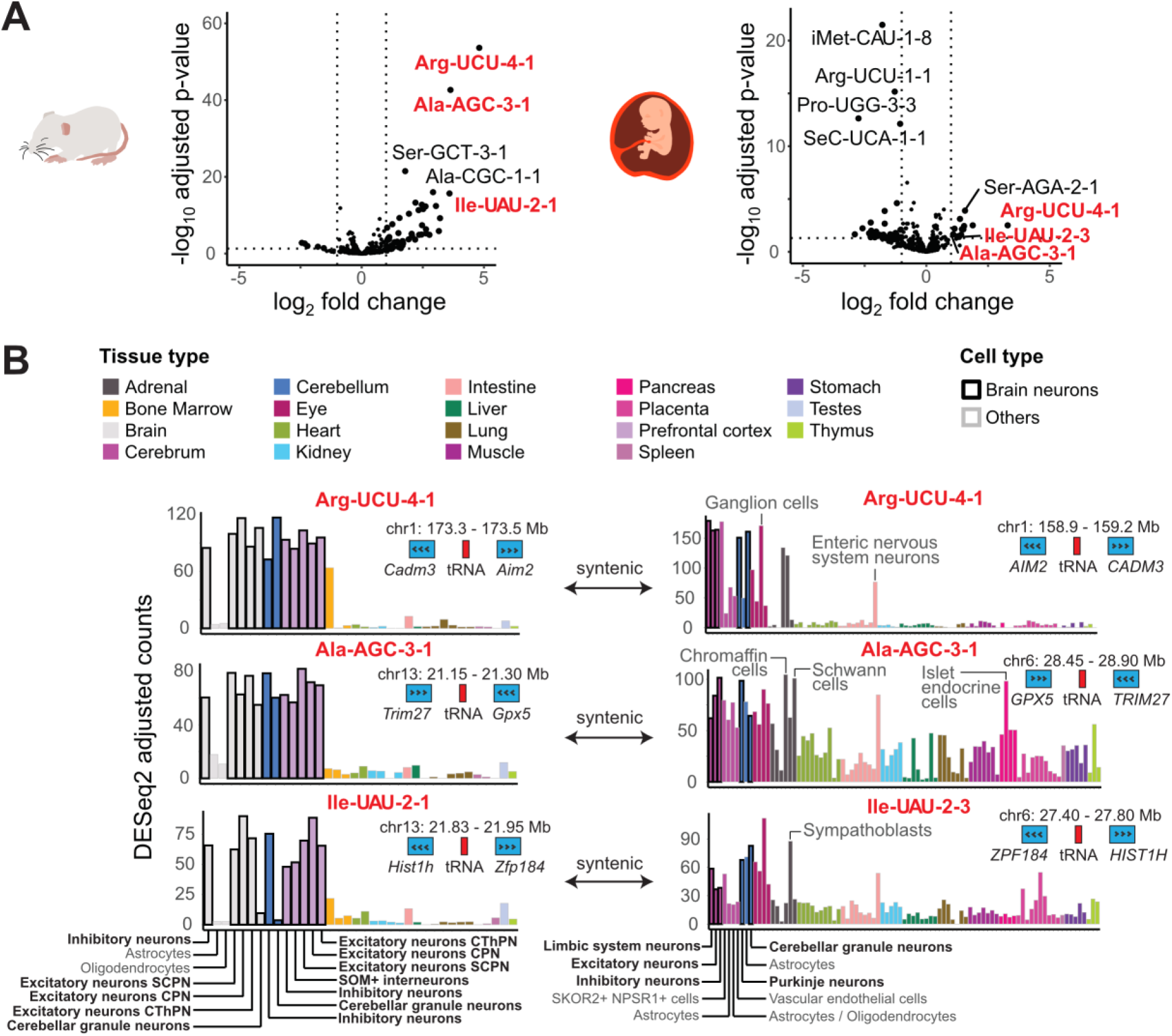
Some tRNA genes are enriched in brain neurons and conserved in mouse and human. **(A)** Volcano plots show individual tRNA gene usage and between brain neurons and all other cell types, displaying −log_10_ adjusted p-values and log_2_ fold change (FC), as determined using DESeq2. Red labels indicate syntenic tRNA genes significantly enriched in brain neurons in both AM (left) and FH (right). Vertical lines indicate a fold change greater than 100% in one direction. **(B)** Bar plots demonstrate expression of syntenic, neuron-enriched tRNA genes. Each bar corresponds to a cell type filled according to the tissue type (far left) and with border shading according as follows: black for brain neurons, gray for others. Brain cell types are labeled at the bottom. In the top right corner of each bar plot is the genomic location of the tRNA gene (red) with flanking protein-coding genes (blue), displayed with arrows denoting strand.

### Amino acid supply and demand are highly correlated across cell types

Having analyzed codon and anticodon usage separately, we then sought to correlate them to assess the potential impact of codon and anticodon pools on translation elongation. Because there is no one-to-one correspondence between codons and anticodons, methods such as the tRNA adaptation index (tAI) have been developed to match multiple cognate anticodons to their respective codons by estimating the stability of various codon-anticodon interactions. With the assumption that highly expressed genes should be most adapted to the anticodon pool, wobbling coefficients can be calculated to measure the correlation between an organism’s codon and anticodon pools (dos Reis et al. 2004). However, synonymous codon usage biases in mammals are not believed to be under strong translational selection from tRNA pools but rather from mutational biases (Dos Reis and Wernisch 2009; Pouyet et al. 2017b). Moreover, we wanted to assess differences in translation efficiency across cell types rather than assume optimal wobble coefficients for translation of the most highly expressed protein-coding genes per cell type. Therefore, we instead measured the correlation between AA demand and AA supply. While this is imperfect as not all tRNAs charged with a particular AA can decode every codon demanding that same AA (Watanabe and Yokobori 2011), this approach allowed us to avoid making assumptions about several variables that have not been quantified at the cell type level, such as the relative decoding efficiency of wobble and Watson-Crick base pairs and the levels of ANN tRNAs that are modified to INN or left unmodified, which should theoretically scale with the expression of adenosine deaminases across cell types.

By defining translation efficiency (TE) as the Spearman’s rank correlation between AA supply and demand from the tRNA and the mRNA side, respectively (**Fig. 6A, Tables S8, S15**), we calculated TE across all AM and FH cell types for which scRNA-seq data were available for AA demand quantification and scATAC-seq data had sufficient resolution for reliable AA supply quantification. We found that TEs range from ρ ~ 0.66 - 0.85 in AM and ρ ~ 0.60 - 0.86 in FH (**Fig. 6A**). This relatively narrow range of TEs is consistent with our observations of stability in both codon and anticodon pools across the majority of cell types, and is also in line with previous bulk analyses using the same TE metric (Kutter et al. 2011).

**Fig. 6:**
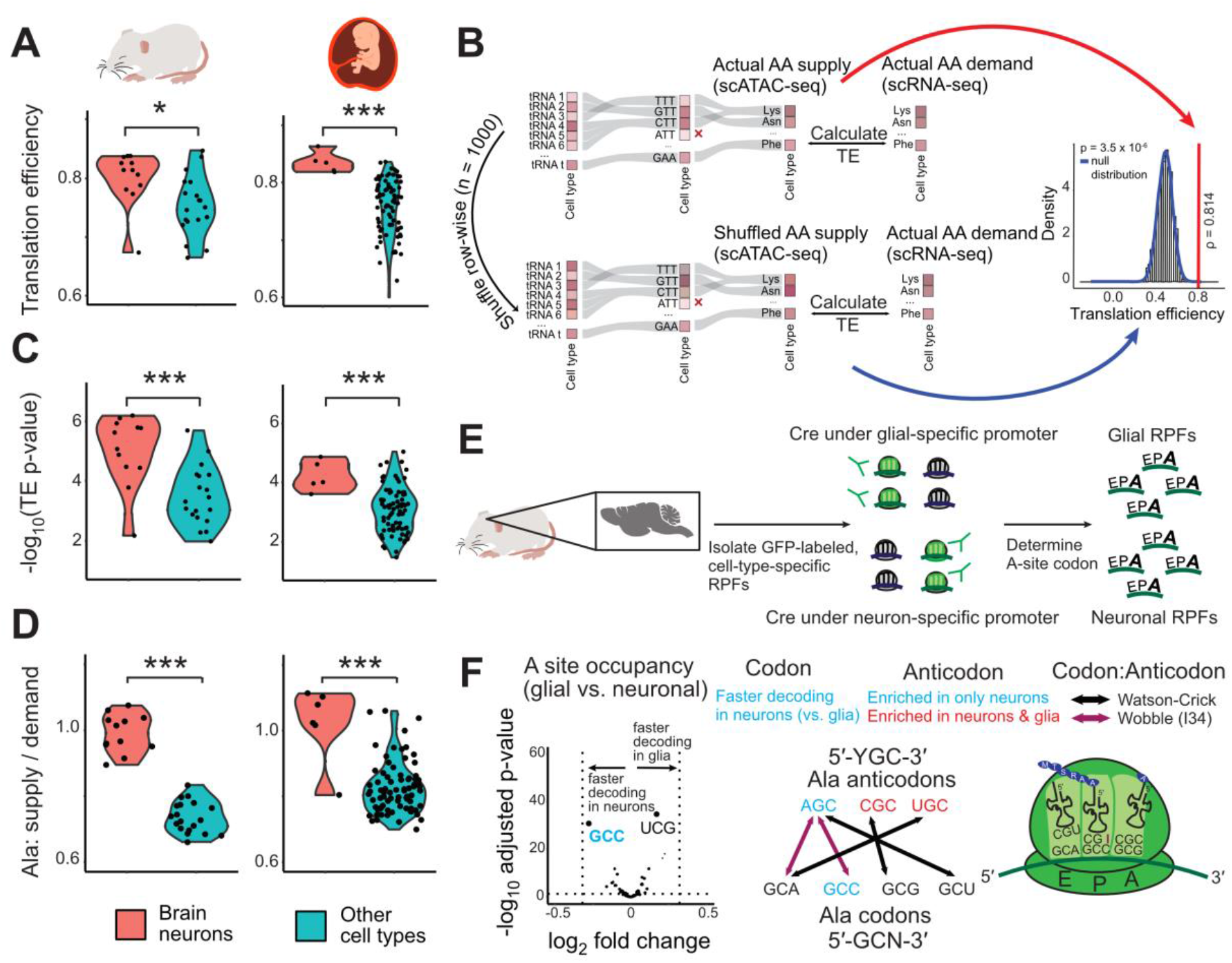
Correlation between amino acid supply and demand is strong and statistically significant in all cell types, with the strongest correlation in brain neurons. **(A, C, D)** Violin plots are divided into AM (left) and FH (right) showing (A) TEs, (C) TE p-values, and (D) alanine (Ala) supply to demand ratio of brain neurons and other cell types. Of all 20 amino acids, only Ala is significantly enriched between brain neurons and all other cell types. (**B**) Schematic representation of the approach used to determine statistical significance of correlation between AA supply (from tRNA side) and AA demand (from mRNA side), defined here as translation efficiency (TE). The observed tRNA expression for each cell type is shuffled 1000 times, pooled at the AA supply level, and correlated to the AA demand (unshuffled) to detect a null distribution of TE values and determine statistical significance of actual TE. (**E)** Workflow from an adult mouse brain ribosome profiling dataset (Scheckel et al. 2020). Cell-type-specific ribosome-protected fragments (RPFs) for neurons and glia were obtained with cell-type specific GFP-labeling of a ribosomal protein, followed by immunoprecipitation against GFP. The ribosome A-site of these cell type-specific RPFs was determined, and differential analysis was performed using DESeq2. **(F)** Volcano plot shows that the tRNA^Ala^ (AGC) anticodon is enriched in neurons and faster decoding of the tRNA^Ala^ (GCC) anticodon is observed in neurons when compared to glial cells. AGC must decode GCC via an adenosine to inosine modification at the first anticodon position. *p<0.05, **p<0.01, ***p<0.001 (Mann-Whitney test).

To assess whether these TEs were statistically significant, we sampled from a null distribution of TEs by shuffling 1000 times the tRNA gene usage of each cell type, pooling the gene expression levels based on each tRNA’s charged AA to calculate AA supply, and then correlating the AA supply to the AA demand from the mRNA side, which was unshuffled (**Fig. 6B**). After estimating parameters for this null distribution, we compared the values of the observed and simulated TEs for each cell type. By using this approach, we discovered that all cell types in AM and all but one cell type (skeletal muscle cells) in FH are statistically significant (p-value < 0.05) (**Fig. 6C**). In other words, codon and anticodon pools were not only stable across different mammalian cell types but also strongly correlated with each other, thereby establishing an efficient interface for translation elongation.

### Neurons have enhanced translation efficiency

While the majority of cell types have similar codon and anticodon pools, some cell types were outliers in either codon usage (cardiac/skeletal muscle cells and pancreatic acinar cells, **Fig. 2**) or anticodon usage (brain neurons, **Fig. 4**) in the AM and FH datasets. We therefore wondered whether the TE varies in these outliers. We found that cardiac/skeletal muscle cells had among the lowest TEs and pancreatic acinar cells had moderate TE when compared other cell types (**Fig. 6A, Tables S8, S15**). While cardiac/skeletal muscle cells differed in codon usage largely because of *TTN*’s immense contribution to their codon pool (**Fig. 2G**), their anticodon pools did not differ. This lack of anticodon compensation for differential codon usage led to the lowest TE in cardiac/skeletal muscle cells. In contrast, we found that brain neurons have among the highest TEs. When comparing brain neurons against all other cell types, we observed that they had a statistically significant increase in both TE and TE p-value (**Fig. 6A, C**). Since we determined that codon usage in neuronal cells was comparable to most other cell types, this increased TE was anticodon-driven.

Because we define TE as the correlation between AA supply and demand, its values are influenced by the similar proportions of each AA’s supply and demand. Therefore, we calculated the ratio of AA supply and demand for each of the standard 20 AAs and compared these supply-demand ratios between brain neurons and other cell types. Ala was the only AA with an AA supply-to-demand ratio that had a statistically significant increase in brain neurons when compared to other cell types (**Figs. 6D, S8**). This is consistent with our finding that Ala anticodons were enriched in brain neurons (**Fig. 4**). Thus, Ala is a prominent driver of increased TE in brain neurons.

### Increased neuronal supply of the Ala-AGC anticodon results in faster decoding of the Ala-GCC codon

Because the Ala supply-demand ratio was enhanced in neurons, we hypothesized it could result in faster decoding of Ala codons. The relative decoding rate can be quantified by ribosome profiling, in which ribonuclease-protected RNAs, including ribosome-protected mRNA fragments (RPFs), are sequenced genome-wide (Ingolia et al. 2019). By mapping RPFs to the genome, codons occupying the ribosome E (exit), P (peptidyl-), and A (aminoacyl-) sites can be determined. Since codon-anticodon recognition takes place within the A-site, codons that are more slowly decoded by the ribosome are expected to more frequently occupy the A-site of RPFs, after adjusting for the background RPF codon frequency. In contrast, faster decoded codons should be less frequent in the A-site of RPFs (Ingolia 2014). Consistent with these expectations, budding yeast in which specific codon-anticodon interactions were disrupted have significantly higher levels of those impaired codons in RPF A-sites when compared to wildtype, suggesting slower decoding (Nedialkova and Leidel 2015).

To test our hypothesis of faster neuronal decoding of Ala codons, we analyzed publicly available cell-type specific ribosome profiling data in mouse brain (Scheckel et al. 2020). Since single-cell ribosome profiling technologies have not yet been developed, Scheckel et al. created mouse cell lines with a GFP-labeled form of a ribosomal protein induced by Cre recombinase under a cell-type-specific promoter. Subsequent affinity purification with an anti-GFP antibody was used to obtain RPFs specific to neuron and glial populations (**Fig. 6E**). After aligning RPFs to the A-sites of protein-coding genes with CONCUR (Frye and Bornelöv 2021), we performed differential analysis between all neuronal and glial samples to examine whether any codons were differentially present in the A-site. Of these codons, Ala-GCC showed the strongest enrichment in glial cells, verifying that neurons decode GCC faster. This is consistent with our analysis of Ala anticodon usage in brain at the single-cell level. We observed that both neurons and glial cells were enriched for Ala-CGC and Ala-UGC when compared to non-brain cell types but were not differentially expressed between glial cells and neurons (**Fig. S3**). In contrast, tRNA^Ala^ (AGC) was specifically enriched in neurons and is the only Ala anticodon that can decode Ala-GCC (**Fig. 6F**). The interaction between the Ala-GCC codon and the tRNA^Ala^ (AGC) anticodon must occur through a A34-to-I modification, in which the first anticodon position A becomes an I that can then base pair with C, the third codon position. While cell-type-specific ribosome profiling datasets are not yet available in other cell types for which we calculated TE, these findings underscore that differential codon and anticodon usage can lead to cell type-specific differences in decoding rates during translation elongation.

Moreover, our identification of an increased tRNA^Ala^ (AGC) anticodon pool in neurons may explain a recent finding regarding the role of *Trm1l*, a tRNA-modification enzyme. Previous studies demonstrated that *Trm1l*-deficient mice had neurological deficits, including altered motor coordination and aberrant exploratory behavior (Vauti et al. 2007). However, the function of this protein was unknown. A recent study described that the TRM1L protein performs N2,N2-dimethylguanosine (m^2,2^G) modifications at position 26 specifically in tRNA^Ala^ (AGC) (Jonkhout et al. 2021). In addition, TRM1L subcellular localization changes upon neuronal activation but not under general stress, suggesting that this protein plays a role in long-term potential and synaptic plasticity. Although it was hypothesized that the neurological phenotype associated with the *Trm1l* deletion could be caused by increased tRNA^Ala^ (AGC) abundance in the brain, differential expression was masked in bulk sequencing data. But with single-cell sequencing, we have now detected this neuronal increase in tRNA^Ala^ (AGC) anticodon usage. Modifications influence the stability of tRNA transcripts and hypomodified tRNA molecules are rapidly degraded (Kimura and Waldor 2019). Thus, absence of the m^2,2^G26 modification upon the deletion of TRM1L may reduce tRNA^Ala^ (AGC) pools in neurons and impact the rate of translation elongation. Future systematic studies may explore whether the absence of tRNA modifications reduces specific anticodon pools and drives malignant phenotypes by slower decoding of the corresponding mRNA codons.

## Discussion

While codon and anticodon usage have been previously quantified in bulk tissue, analysis of these critical players in translation elongation had not yet been explored at single-cell resolution. By harnessing publicly available scRNA-seq and scATAC-seq atlases, we simultaneously analyzed codon usage and AA demand as well as anticodon usage and AA supply across cell types from multiple tissues in two mammals. The highly comprehensive nature of these atlases allowed us to examine codon and anticodon usage not only in greater depth but also breadth since the AM and FH atlases contain more tissues than any bulk dataset.

Since existing tRNA quantification methods cannot yet be performed at the single-cell level, we investigated the feasibility of using scATAC-seq and demonstrate that it is robust for measuring tRNA gene usage. The increasingly lower cost and proliferation of many fast and reproducible scATAC-seq methods may make this approach especially attractive for tRNA quantification. In particular, scATAC-seq breaks a major bottleneck that has hindered comprehensive analysis of tRNAs across the diverse cell types in complex mammalian systems. While our analysis of codon and anticodon usage data may have biases arising from integrating data from two different techniques (scRNA-seq and scATAC-seq) performed on different samples, the increased proliferation of multi-omics techniques that can jointly quantify chromatin accessibility and transcripts in single cells should remove this limitation (Swanson et al. 2021). Another issue with scATAC-seq is that few cuts to tRNA genes are sequenced under normal sequence depths, which necessitated removing some cell types with insufficient cuts from our analysis (**Fig. 3C**). However, a multi-omics approach that combines scRNA-seq (for cell type annotation) and single-cell CUT&Tag (Kaya-Okur et al. 2019) to enrich specifically for Tn5 insertions near Pol III binding sites should mitigate this problem.

While we identify here several striking features of codon usage, anticodon usage, and translation efficiency that are conserved over 90 million years of mammalian evolution (Kumar et al.), we found an increase of Ala supply in AM and FH brain across all datasets (bulk and single-cell). Besides Ala, other AA are enriched in some brain datasets but not in others. As a result, the apparent faster decoding of the codon Cys-UCG in glial cells within the ribosome profiling dataset remains to be resolved (**Fig. 6F**). AA supply and demand are known to change at timescales of minutes and hours. These fluctuations across datasets may reflect differences right before sample collection (Rak et al. 2018). In contrast, the evolutionarily conserved nature of increased tRNA^Ala^ (AGC) supply in both mammals at different time points suggests purifying selection and a longer-term significance in organismal development.

Another potential area for progress involves better modelling the rate of translation elongation. When calculating TE, we determined AA supply from the usage of all tRNA genes charged with the same AA and calculated AA demand by weighting the AA gene usages by their expression for all cell types. From the AA supply side, it is oversimplistic to assume that all tRNAs will necessarily function in translation elongation, especially since tRNAs are known to perform an array of different roles, including as tRNA-derived fragments (Polacek and Ivanov 2020). From the AA demand side, different protein isoforms are known to exist across cell types. We have not accounted for this due to the lack of data availability, although the stability of codon usage across cell types that express entirely different proteins may indicate that including isoform data will not affect our main conclusions. For both AA supply and demand, it is important to not only consider the abundances of tRNAs and mRNAs but also their turnover rates. Once single-cell half-life information of mRNA and tRNA species become feasible to measure, AA supply and demand should account not only expression but also the differential stability of these transcripts. Additionally, codon optimality has been shown to be a major determinant of mRNA stability (Presnyak et al. 2015). Thus, the ratios of corresponding codons and anticodons or AA supply and demand across different cell types could be correlated with the stability of mRNA transcripts. In particular, it would be interesting to see if neuron depletion of tRNA^Ala^ (AGC) anticodon results in slower decoding of Ala-GCC codons, inducing slowness-mediated decay of certain transcripts vital for neuronal function (Rak et al. 2018).

For translation efficiency, we calculated TE at the level of AA supply and demand, rather than at the codon-anticodon level as it is unclear whether and how to consider wobble base pairing and other modulators of the codon-anticodon interaction such as tRNA modification enzymes. Moreover, there may be tissue-specific tRNA modifications, including a recent finding that some tRNA^Ala^ anticodons are enriched for particular modifications in the brain (Pinkard et al. 2020). Thus, another avenue of investigation will involve disentangling cell type differences in these processes that could impact the rate of translation elongation.

Finally, it is worth noting that our analyses were performed on atlases of mice and human samples believed to represent healthy states. Thus, the stability that we observe in codon and anticodon pools may exist only in a healthy state, and dysregulation of these pools may occur in abnormal states such as cancer (Goodarzi et al. 2016; Zhang et al. 2018), in neurodegenerative diseases (Kapur et al. 2017), and with disruption to the microbiome (Schwartz et al. 2018; Huang et al. 2021). Though such single-cell data are not yet available, approaches similar to those presented here could be used to examine cell-type-specific changes to codon usage, anticodon usage, and translation efficiency in disease.

## Materials and Methods

### Datasets analyzed and code availability

All data analyzed were downloaded from publicly available sources (refer to Table S1 for more information). The adult mouse (AM) scRNA-seq atlas was obtained from GSE109774 (Ingolia et al. 2019) and Seurat objects were downloaded from the Tabula Muris website (https://tabula-muris.ds.czbiohub.org/). The AM scATAC-seq atlas (Schaum et al. 2018) was obtained from GSE111586 and BAM files were downloaded from the website (https://atlas.gs.washington.edu/mouse-atac/). The fetal human (FH) scRNA-seq (Cusanovich et al. 2018) and FH scATAC-seq atlases (Cao et al. 2020) are available from GSE156793 and GSE149683, respectively. Count matrices (scRNA-seq) and fragment files (scATAC-seq) dataset were downloaded from the website (https://descartes.brotmanbaty.org/bbi/). Two other mouse brain scATAC-seq datasets were analyzed to determine scATAC-seq reproducibility (**Figs. 3C, S2**): a 10X Genomics dataset (atac v1 adult brain fresh 5k) and a droplet-based dataset from Lareau et al. (GSE123581) (Lareau et al. 2019). scATAC-seq tRNA gene usage was also compared with bulk adult mouse ATAC-seq data (SRA: SRX4946150) (Liu et al. 2019) and ChIP-seq data (ArrayExpress: E-MTAB-2326) (Schmitt et al. 2014). To compare with anticodon usage results from scATAC-seq concerning increased alanine supply, bulk experiments were analyzed. A bulk ATAC-seq atlas of early mouse development was downloaded from https://www.encodeproject.org/ (Gorkin et al. 2020). Bulk ChIP-seq data of mouse liver and brain across development were downloaded from Array Express accession E-MTAB-2326 (Schmitt et al. 2014). QuantM-tRNA-seq of adult mouse tissues was downloaded from GSE141436 (Pinkard et al. 2020). To examine differential decoding of codons in neurons versus glial cells, the cell-type specific mouse brain ribosomal profiling dataset (Domcke et al. 2020) was downloaded from GSE149805.

The code, written in R, is available on Github at https://github.com/wgao688/sc_tRNA_mRNA. Tidyverse was used for data exploration and analysis, and ggplot was used for data visualization. Several packages in the Bioconductor suite were used (Gentleman et al. 2004). Seurat was used for scRNA-seq analysis (Satija et al. 2015), and Signac was used for scATAC-seq analysis (Stuart et al.). DESeq2 was used for differential gene analysis (Love et al. 2014). tRNAscanImport (https://github.com/FelixErnst/tRNAscanImport) was used to load tRNA gene predictions for mm10 and hg19. Gviz was used for genome browser visualization (Hahne and Ivanek 2016).

Count matrices, pooled at the cell-type level, for mRNA gene expression, codon usage, and AA demand, as well as for tRNA gene expression, anticodon usage, and AA supply are available as Supplemental Data (Tables S2-S7 and S9-S14 for AM and FH, respectively).

### Quantification of codon usage and amino acid demand from scRNA-seq datasets

Codon usage for each protein-coding gene was determined using the Ensembl set of protein-coding genes for mouse (*Mus musculus*, mm10) and human (*Homo sapiens*, hg19). This was then formatted as a codon frequency per gene matrix of dimensions 61 × *p*, where the rows correspond to the 61 sense codons and the columns to each of the *p* protein-coding genes. Codon usage for each individual cell was determined by matrix-multiplying the codon frequency per gene matrix by the scRNA-seq count matrix, formatted as a *p* × *n* matrix where *p* again corresponds to the protein-coding genes (ordered in the same way as the codon usage matrix) and *n* to the number of cells remaining after quality control filtering. Only genes present in both the count matrix and the codon usage were used, since *p* must be the same for matrix multiplication. The multiplication of these two matrices weights the codon frequencies of each protein-coding gene by the expression in each cell, producing a “codon usage per cell matrix” of dimension 61 × *n*.

Using the cell type annotations provided by the atlases used in this study, we can pool the codon usage of cells of the same cell type together, producing a “codon usage per cell type” matrix of dimension 61 × *m*, where *m* denotes the number of cell types annotated in the experiment. Each codon has a specific amino acid demanded per the standard genetic code. Therefore, amino acid demanded can be quantified by pooling the codons demanding the same amino acid, to produce an “amino acid per cell type” matrix of dimension 20 × *m*, where the rows correspond to the twenty classical amino acids.

### Quantification of tRNA gene usage, anticodon usage, and AA supply from scATAC-seq datasets

tRNA gene annotations were used from gtRNAdb (Chan and Lowe 2016) that uses tRNAscan-SE v2.0 to predict tRNA genes. tRNAscan-SE denotes functional tRNAs as “high confident” based on predicted secondary structure stability and other tRNA-specific features whereas “low confident” are considered as probable pseudogenes. There are 401 high-confidence tRNA genes in adult mouse (mm10 annotation) and 416 in human (hg19 annotation). Quantification of tRNA gene usage from scATAC-seq datasets required generating a tRNA count matrix of dimension *t* × *n*, where *t* corresponds to the number of high-confidence tRNA genes. For scATAC-seq datasets, this quantification was performed using the FeatureMatrix function available in Signac, which uses a fragment file and a set of genomic regions as input (Stuart et al.). Fragment files were already available for the human fetal dataset. For the AM dataset, BAM files were downloaded from https://atlas.gs.washington.edu/mouse-atac/and sinto (https://timoast.github.io/sinto/) was used for generation of fragment files. FeatureMatrix quantifies the number of Tn5 insertions occurring within each of the specified genomic regions (formatted as a Granges object). In our case, the genomic regions were the set of high confidence tRNA genes, including the gene body and 100 nucleotides upstream and downstream, as has been performed in many other studies using Pol III ChIP-seq (Kutter et al. 2011).

This “tRNA gene expression per cell” matrix can be pooled by cell type annotation to produce a “tRNA gene expression per cell type” matrix of dimension *t* × *m*, where m refers to the number of cell types annotated. This matrix was then used to examine the total number of cuts for each tRNA gene across all cells (pseudobulked), and a bimodal distribution was observed, interpreted as background and true signal distribution. tRNA genes belonging to the background distribution were removed from the analysis. tRNA genes can then be pooled based on their anticodon sequences to produce an “anticodon usage per cell type” matrix of dimension *a* × *m*, where *a* refers the number of unique anticodons (46 in mm10 and 47 in hg19). Finally, amino acid supply can be quantified by pooling anticodon families by the amino acid they accept, resulting in an “AA supply per cell type” matrix of dimension 21 × *m*. The 21^st^ amino acid corresponds to selenocysteine.

### Differential analysis

Differential tRNA gene expression, anticodon usage, AA supply analysis was performed using DESeq2 (Love et al. 2014) under default settings, with input being the tRNA gene, anticodon usage, and AA supply per cell type matrices, respectively. Brain neurons were compared against all other cell types in the adult mouse and fetal human datasets.

### Translation efficiency analysis

We compute translation efficiency (TE) as the Spearman’s rank correlation coefficients between the amino acid demand from the mRNA codon side and the amino acid supply from the tRNA anticodon side. In other words, the values for each cell type in the “AA demand per cell type” matrix and the “AA supply per cell type” matrix were correlated with each other, for all cell types that were present in both the scRNA-seq and scATAC-seq datasets. For this analysis, selenocysteine was ignored since the number of stop codons that bind to the selenocysteine tRNA are unknown at the cell type level.

### Ribosome profiling analysis

Scheckel et al. performed ribosome profiling on mice injected with control homogenate or prions and extracted cell-type specific ribosome-protected fragments pertaining to CamKIIa excitatory neurons, PV interneurons, microglia, and astrocytes (Scheckel et al. 2020). They collected samples from mice sacrificed 2, 4-, 8-, 16-, and 24-weeks post injection, as well as samples collected at a later point before severe prion disease developed in the prion-infected group. We downloaded the ribosome profiling data for all time points for only the control groups. Fewer ribosome-protected fragments (RPFs) at the A-site, when corrected for background codon frequencies at other positions in RPFs, suggests faster decoding of a particular codon. To identify A-site codon for each RPF, we used CONCUR (Frye and Bornelöv 2021). To identify codons that were differentially found in the ribosome A-site, we used DESeq2 (Love et al. 2014), as described above, to compare the neuronal group (CamKIIa excitatory neuron and PV interneuron samples at all time points) and the glial group (microglia and astrocyte cell types), while controlling for injection time.

## List of abbreviations

RNA: ribonucleic acid
tRNA: transfer RNA
mRNA: messenger RNA
sc: single-cell
RNA-seq: RNA-sequencing
ATAC-seq: Assay for Transposase Accessible Chromatin-sequencing
AA: amino acid
ChIP-seq: Chromatin Immunoprecipitation with massively parallel DNA sequencing
RP: ribosome profiling

## Data access

No new data was generated for this study, as all data analyzed in this paper were downloaded from publicly available datasets. The sources of these datasets are detailed in the pertinent sections of the Methods.

## Competing interests

The authors declare no competing financial and non-financial interests.

## Author contributions

WG and CK conceptualized the project. WG, CG, and CK designed the analyses. WG performed the analyses. WG, CG, and CK wrote the original draft.

## Acknowledgements

We highly appreciate critical comments on the manuscript by Konrad Rudolph. We thank Keyi Geng and Marcel Tarbier as well as other group members of the laboratories of Claudia Kutter, Marc Friedländer and Vicent Pelechano for helpful feedback regarding the experimental procedures, data analysis, and data presentation. We are grateful to the Tabula Muris and Descartes consortia for sharing invaluable data to the research committee.

## Financial Support

This work was supported by the U.S.-Sweden Fulbright Student Research Program (WG), Knut & Alice Wallenberg foundation (KAW 2016.0174, CK), Ruth & Richard Julin foundation (2017–00358, 2018–00328, 2020-00294 CK), SFO-SciLifeLab fellowship (SFO_004, CK), Swedish Research Council (2019–05165, CK), Lillian Sagen & Curt Ericsson research foundation (2021-00427, CK), Gösta Miltons research foundation (2021-00527, CK), and the Swedish National Infrastructure for Computing projects at UPPMAX (storage: 2021/22-133 and compute: 2021/22-133).

## Notes

### Competing Interest Statement

The authors have declared no competing interest.

